# Naturalistic affective behaviors decoded from spectro-spatial features of multi-day human intracranial recordings

**DOI:** 10.1101/2020.11.26.400374

**Authors:** Maryam Bijanzadeh, Ankit N. Khambhati, Maansi Desai, Deanna L. Wallace, Alia Shafi, Heather E. Dawes, Virginia E. Sturm, Edward F. Chang

## Abstract

Task-based studies have uncovered distributed neural networks that support emotions, but little is known about how these networks produce affective behaviors in non-laboratory, ecological settings. We obtained continuous intracranial electroencephalography (iEEG) recordings from the emotion network in 11 patients with epilepsy during multi-day hospitalizations. We coded naturalistic affective behaviors (spontaneous expressions of positive or negative affect) from 116 hours of time-locked video recordings obtained over multiple days from subjects’ hospital rooms and utilized data driven classifiers to determine whether we could decode naturalistic affective behaviors from the neural data. Results indicated that binary within-subject random forest models could decode positive and negative affective behaviors from affectless behaviors (behaviors lacking valence) with up to 93% accuracy. Across the emotion network, positive and negative affective behaviors were associated with increased high frequency activity and decreased lower frequency activity. The anterior insula, amygdala, hippocampus, and anterior cingulate cortex (ACC) made strong contributions to affective behaviors in general. In a subset of subjects, three-state decoders distinguished among the positive, negative, and affectless behaviors using the spectro-spatial features from the emotion network. This study demonstrates that multi-day, highly resolved iEEG recordings in cortical and deep brain structures can reveal the circuit-level physiology of affective behaviors. By measuring behavior in an ecologically valid setting, our findings provide novel insights into the spatially distributed dynamics of local neural populations underlying naturalistic affective behaviors.

**Significance:** Previous neuroimaging and neurophysiological studies have identified a distributed network that supports emotions, but much remains unknown about how this network produces affective behaviors in ecological settings. We used intracranial electroencephalography recordings from the emotion network to decode naturalistic affective behaviors—spontaneous expressions of positive and negative affect that occurred during multi-day hospital stays—from neural data in patients with epilepsy. Our results complement prior neuroscientific studies of emotion and offer novel insights into the spectral and spatial dynamics of the emotion network that characterize naturalistic affective behaviors. The present study suggests intracranial electroencephalography can uncover new details about emotion network physiology and help to expand current neuroanatomical frameworks of human emotions.

## Introduction

Emotions arise in everyday life but are primarily studied in task-based paradigms with limited personal relevance. In the real world, emotions can be powerful, shaping day-to-day decision-making and behavior and fostering social communication and relationships(1). How the brain produces emotions on a day-to-day basis is largely unexplored due to the methodological constraints of functional magnetic imaging (fMRI) studies, which typically involve subjects passively perceiving or actively rating emotionally evocative stimuli while lying in a scanner. Although fMRI studies have provided a wealth of information about the neural networks that support emotion perception, generation, and regulation, they are not able to measure rapid changes in emotion network electrophysiology or spectral activity in different frequency bands, relatively unexplored facets of emotions that may help to reveal their neuroanatomical basis. By providing direct recordings of neural activity in deep brain structures on a millisecond level resolution (2, 3)., intracranial electroencephalography (iEEG) offers a novel window into the neural mechanisms underlying emotions and the behaviors accompanying them.

Neuroimaging studies have elucidated a distributed brain network that is critical for the generation of emotions. In our view, emotions are brief states accompanied by coordinated, widespread changes across the body (e.g., in the face, voice, somatic muscles, and autonomic nervous system) that serve specific survival-relevant functions. While some previous fMRI studies have emphasized the roles of certain regions in different emotions—such as the insula in disgust (4–8), subgenual anterior cingulate cortex in sadness, amygdala in fear (9, 10), and ventral striatum in joy (11), there is also substantial evidence that the emotion network largely responds in a similar way across states of even different valence and arousal (12). Numerous fMRI studies have shown that during emotions there is elevated activity in an emotion-relevant salience network, a distributed network anchored by the anterior insula and anterior cingulate cortex (ACC) (4–8, 13). The anterior insula and ACC have tight reciprocal connections and, through direct projections to subcortical central pattern generators (e.g., amygdala), are critical for producing and sensing visceromotor changes, including those that arise during emotions, in the internal milieu via interoceptive pathways (7, 14–20). Activity in this network increases as emotions intensify but decreases with engagement of emotion regulation systems. With hubs in lateral orbitofrontal cortex, frontoparietal networks are critical for modulating emotions to meet prevailing goals and for reducing activity in emotion generating structures such as the amygdala.

Non-invasive, scalp-based electroencephalography (EEG) studies in humans and local field potential studies in animals have also helped to delineate how the emotion network operates. EEG studies have demonstrated that structures within the emotion network exhibit rapid responses in local field potentials to affective stimuli. For example, the insula responds to faces expressing disgust within 300 ms (9, 21–25), and centro-parietal sites respond to pleasant and unpleasant images within 250 ms (26). EEG studies have also offered some evidence that different patterns of activity and connectivity in specific frequency bands across the emotion network may characterize different affective states. Images that elicit negative emotions, for example, are associated with increased activity in the high gamma band in some structures more than others (e.g., amygdala) (9) and with altered connectivity between certain regions when measured in specific frequency bands (27). Although few EEG studies have compared network activity during different affective states, those that have included multiple emotion conditions have found mixed results as to whether different spectral changes distinguish between positive and negative emotions (28–30). While some task-based studies have found high gamma band activity in the emotion network increases during both positive (28, 31, 32) and negative emotions (9, 28, 30, 33, 34), others have found that activity in lower frequency bands (e.g., theta and alpha) increases during positive emotions (35). Whether fast and slow neural oscillatory changes accompany naturalistic emotions is not well understood, but a more complete mapping of emotion network activity across multiple frequency bands may help to uncover different neural patterns during different affective states.

Here, we examined whether direct iEEG recordings from the emotion network could be used to decode real-world positive and negative affective behaviors from behaviors lacking affect (“affectless behaviors”). iEEG studies of emotion are few but offer certain advantages over other techniques. In addition to sampling deep brain structures, iEEG enables recording of neural activity at higher temporal resolutions and with greater spatial specificity than is possible through non-invasive imaging techniques. iEEG recordings can also span longer periods of time than fMRI and EEG studies and are not limited by the task-based paradigms. iEEG is sensitive to high frequency changes with greater signal-to-noise ratio than scalp EEG and enables study of fast and slow neuronal processes such as high (e.g., high gamma) and low (e.g., theta) frequency activities. Although little is known about the role of different frequency bands in everyday affective experience, recent iEEG studies have also found that lower mood is associated with greater activity in low frequency bands (e.g., beta) (36) and that deep brain stimulation of lateral orbitofrontal cortex can decrease low frequency (i.e., theta band) activity in the emotion network and improve mood (37).

Previous studies have used iEEG to decode speech(38), everyday behaviors (e.g., television watching)(39), and self-reported mood (40) from iEEG recordings, but whether decoders can successfully discriminate positive and negative affective behaviors from affectless behaviors using neural recordings has not yet been investigated. During a multiple day hospital stay, we obtained continuous, multi-site iEEG recordings of the human emotion network in individuals undergoing surgery for intractable epilepsy. Electrode placement was based on the clinical needs of each person and, thus, varied across people but covered numerous sites within the emotion network, including anterior insula, anterior cingulate cortex, orbitofrontal cortex (OFC), amygdala and hippocampus. Using video recordings of the participants in their hospital rooms, we annotated the affective behaviors they displayed spontaneously throughout their stay (herein, “naturalistic affective behaviors”) and then examined simultaneous iEEG neural recordings during those moments.

We first tested whether the decoders could differentiate among positive, negative, and affectless behaviors, and then examined the features of emotion network activity that influenced their performance. Although fMRI and EEG studies have suggested that similar changes across the emotion network characterize a number of affective states, how the emotion network produces these different states is not well understood. Changes in power in specific frequency bands (i.e., the spectral features) in electrodes located in specific network hubs (i.e., the spatial features) could together create unique “spectro-spatial” patterns across the emotion network that accompany different behaviors. Our central hypotheses were that common spectral changes across the emotion network might characterize both positive and negative affective behaviors but that each behavior would also be characterized by its own unique spectro-spatial features.

Positive and negative affective behaviors, by definition, differ in valence, but they can also have similar qualities such as comparable levels of arousal (i.e., intensity), which may be represented by common changes across the emotion network. Given that gamma band activity is characterized by fast oscillations thought to reflect neuronal activity in humans (41), we expected that both positive and negative affective behaviors would be characterized by increased gamma activity in the emotion network compared to affectless behaviors. We also hypothesized that gamma activity in the anterior insula and ACC, regions important for interoception and emotion generation(19, 42), would contribute more strongly to the production of both types of affective behaviors than OFC, a region in which lateral areas support emotion regulation and behavioral control. We also investigated lower frequency bands and tested whether increased low frequency activity was associated with negative affective behaviors, as stimulation studies of mood would suggest, or whether it was a feature of positive and negative states more generally.

Although distributed changes in spectral activity across the emotion network may characterize positive and negative affective behaviors in general, we expected that spatial differences might also differentiate among the behaviors as some regions might play more prominent roles in certain behaviors more than others. When stimulated, the ventral ACC can induce positive behaviors and feelings such as laughter and mirth (16, 18, 33, 43), and the dorsal ACC can produce feelings of impending doom and perseverance. The amygdala is primarily known for its predominant role in negative affect but can be active during positive states of sufficient intensity. We expected that ventral ACC might contribute more strongly to positive affective behaviors and that the dorsal ACC and amygdala would contribute more strongly to negative affective behaviors (21,24, 31).

## Results

We obtained 24-hour bedside audiovisual recordings and continuous iEEG data in subjects with intractable epilepsy who were hospitalized for clinical seizure monitoring with temporary subdural electrodes (Fig. 1-A, Appendix, Table S1). To examine the neural mechanisms underlying naturalistic affective behaviors, we analyzed a total of 116 hours (mean = 10.5 hour, SD = 5.48) of behavioral (Fig 1-B) and neural data (Fig 1-C) in 11 subjects with electrodes placed in at least three mesolimbic structures, which here included the anterior cingulate cortex (ACC), anterior insula, OFC, amygdala, and hippocampus (Appendix, Table S1).

**Figure 1.**
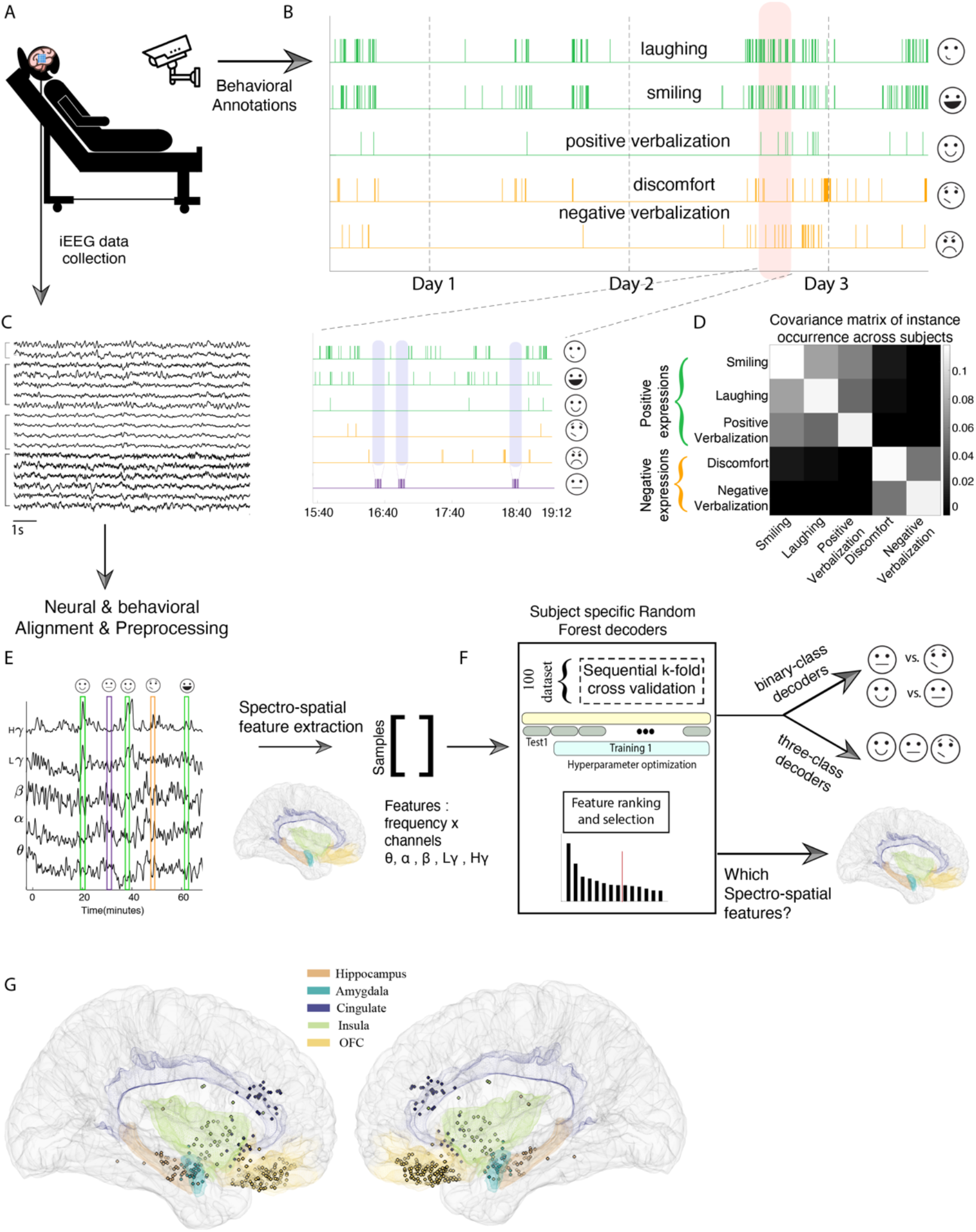
Collection and processing pipelines for the behavioral and neural data streams. A) Schematic of an example patient during the multi-day hospital stay who undergoes continuous measurement of neural and video recordings. B) Video recordings were hand annotated to identify instances of positive affective behaviors (green), negative affective behaviors (orange), and affectless behaviors. In the inset, we zoom in on a three-hour period (orange shading) to illustrate examples of affectless behaviors (purple shading). C) 10 seconds of raw iEEG data traces from four regions are provided as an example. D) Covariance matrix of occurrences of affective behaviors across subjects. E) Magnitude of Hilbert transform in five frequency bands overlaid with instances of affective behaviors for an example insula channel across 60 minutes. F) The pipeline for training the random forest decoder models. G) Right and left view of MNI brain, showing the electrode coverage within emotion network structures across all subjects.

A team of human raters (11 total) blind to the study’s goals and hypotheses later hand-annotated the video recordings to denote the subjects’ spontaneous behaviors (Fig.1-B & Appendix, Table S2, Fig. S1-A). Raters used the linguistic annotation software ELAN (44) to mark various behaviors such as conversation, drinking, smiling, etc. (Appendix, Table S2). We focused our analyses on the affective behaviors, which included instances of smiling, laughing, positive verbalizations (i.e. when the subject spontaneously expressed positive feelings such as, “I’m happy!”), pain-discomfort, and negative verbalizations (Fig.1-B, Appendix, Table S1). We defined “affectless behaviors” as neutral 10-minute periods in which there were neither positive nor negative affective behaviors in the audiovisual recordings (purple shading in Fig.1-B, bottom panel). These periods were often characterized by other naturalistic behaviors such as engagement in conversation that lacked clear affect.

To characterize the associations among different affective behaviors, we counted the number of occurrences of the affective behaviors (i.e., smiling, laughing, positive verbalizations, negative verbalization, and pain-discomfort) and found that smiling, laughing, positive verbalizations positively covaried. For example, subjects who displayed more of one behavior (e.g., smiling) were also more likely to display more of another (e.g., laughter). Thus, we grouped these behaviors into a single “positive affective behaviors” category. Pain-discomfort and negative verbalizations also positively covaried (Fig. 1-D) and were grouped into a “negative affective behaviors” category. Overall, our dataset included more instances of positive behaviors (mean ± sem = 112 ± 17, n = 10 subjects) than negative behaviors (mean ± sem = 61 ± 19, n = 5 subjects). While smiling and laughing occurred frequently, pain-discomfort and negative verbalizations were less common (Appendix, Fig. S1-B). A subset of videos was annotated by second raters to assess inter-rater reliability, and overall there was high agreement between the raters. 82% of the total instances of affective behaviors that were recorded by one rater were also recorded by the other. The instances were highly overlapping in time with the onset of each instance having a median difference of 0.87 seconds (mean = 7.4 sec, Appendix, Fig. S1-C) between any two raters.

When examined in relation to the neural data, subjects exhibited a wide range of positive (range: 42-164), negative (range: 34-133), and affectless (range: 277-499, Appendix, Table S3, e.g. Subject 3) behaviors that were aligned with clean neural signals free from epileptic activity (Appendix and Fig. S2). Although subjects had extensive electrode coverage within mesolimbic regions (Fig. 1-G), electrode placement was based on each subject’s clinical needs and, thus, varied across individuals. In each subject, we extracted the spectral power in conventional EEG frequency bands (i.e., theta: 4-8 Hz, alpha: 8-12 Hz, beta: 12-30 Hz, low gamma: 30-55 Hz, and high gamma: 70-150Hz) from electrodes in mesolimbic structures. We computed the average power in each frequency band (i.e., the spectral features) for each electrode (i.e., the spatial features) using 10-second non-overlapping window centered on each positive, negative, or affectless behavior. Together, we refer to these as “spectro-spatial features” as each feature reflected activity in a specific region in the network in a specific frequency band.

We used an objective approach to examine the spectro-spatial features that characterized naturalistic affective behaviors. Briefly, we formed matrices of spectro-spatial features to train binary and multi-class decoder models (Fig.1 E-F). The goal of the binary decoder was to decode affective (either positive or negative) from affectless behaviors, and the goal of the multi-class decoder was to decode positive, negative affective and affectless behaviors. We first trained the decoders in individual subjects and then looked across the group for common findings across the sample. We also conducted detailed investigations to determine how spectral and spatial activity patterns across the mesolimbic network contributed to the decoders’ performance and ability to discriminate among affective states.

### Within-subject random forest models decoded positive and negative affective behaviors from affectless behaviors with up to 93% accuracy

We first trained binary decoders to determine whether we could distinguish affective behaviors from affectless behaviors in each subject separately. The goal of the *positive decoder* was to distinguish positive affective behaviors from affectless behaviors (n = 10 subjects), and the goal of the *negative decoder* was to distinguish negative affective behaviors from affectless behaviors (n = 5 subjects). We constructed random forest (RF) models, which are ensembles of decision trees, and trained them on the spectro-spatial features for the positive, negative, and affectless behaviors (see Figure 2-A for an example subject).

**Figure 2.**
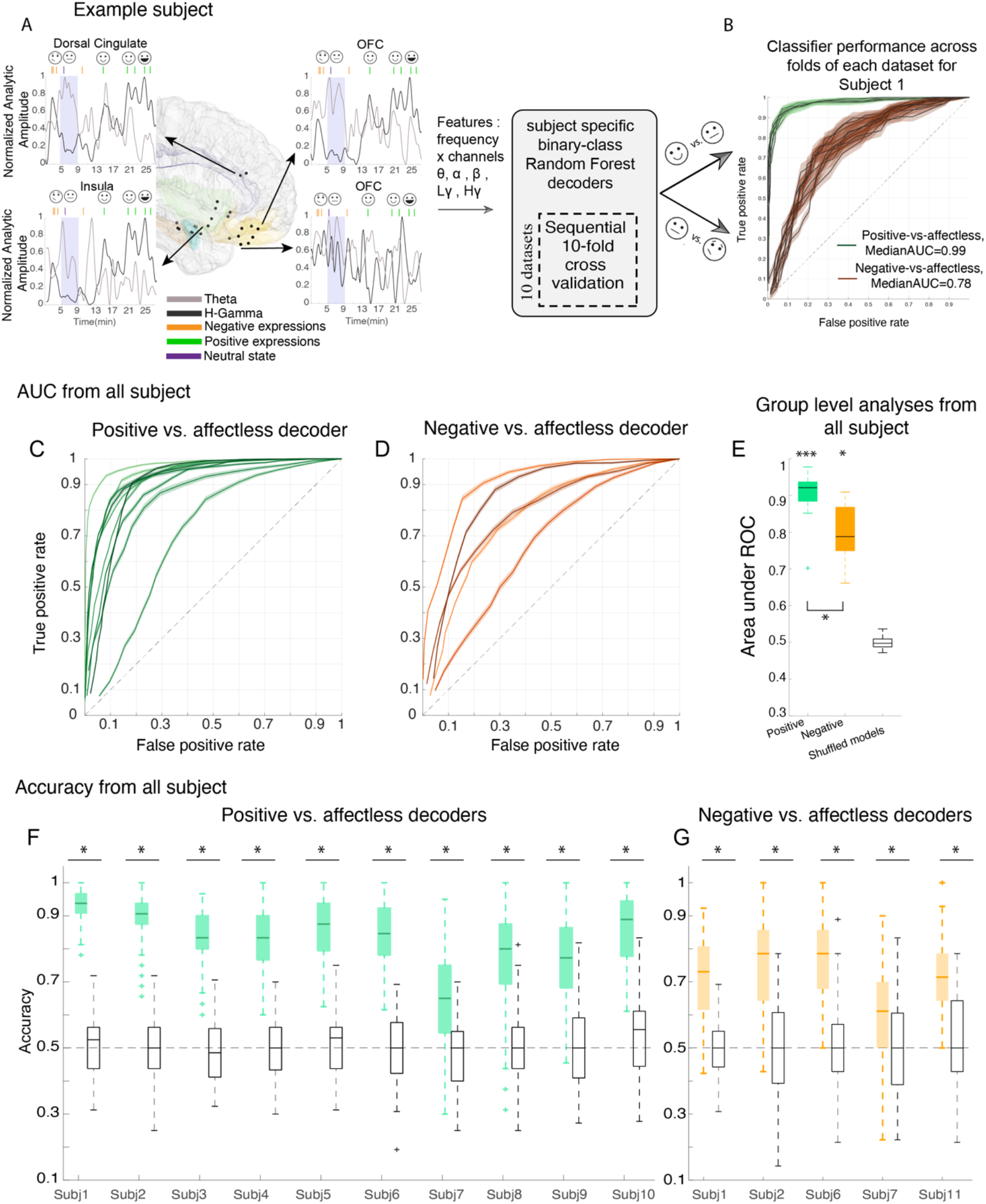
Within-subject random forest models decode positive and negative affective behaviors from affectless behaviors. A) The MNI brain is shown for an example subject to illustrate the locations of the leads (black dots) and measured spectral features that were used in the decoder models. Insets indicate 27 minutes of high gamma (black) and theta (gray) analytic amplitudes for four example channels that are aligned with the affective behaviors in green and orange for the example subject; purple shadings shows affectless periods. B) ROC curve for the example subject across 10 datasets (in which affectless states are selected from different recording times to reduce selection bias; see Appendix) for positive decoders (green) and negative decoders (orange). The shadings represent the SEM across 10 folds. C & D) Area under ROC curve (AUC) for all 10 and 5 subjects that positive & negative decoders were trained on, respectively. Each solid line represents one subject. The shadings are SEM across all datasets. E) Distribution of AUC for positive (green), negative (orange), and permuted models (black) that were trained the same way using the shuffled labels across all subjects. F & G) Accuracies of the same models as in C and D.

We next organized the spectro-spatial features into a balanced dataset (with equal number of instances from each class of behavior) to mitigate sampling bias. As behavioral instances could have occurred close in time, which could lead to artificially high correlations between neural features and model overfitting, we used a conservative sequential k-fold cross-validation classifier to train the RF models (refer to Appendix). To evaluate the decoding performance of our models, we then constructed surrogate permutation models by shuffling class labels (i.e., positive and affectless labels were shuffled within each fold, refer to Appendix). We found the positive and negative decoders could discriminate affective behaviors from the affectless state significantly better than chance in each subject (50%). For example, the median area under receiver operating characteristic (ROC) curve (AUC) across the folds of positive and negative behaviors for the example subject are 0.98 and 0.82, respectively (Fig. 2-B, Subject 1).

To investigate whether these within-subject results were robust and evident across individuals, we next trained the RF classifiers in all subjects. This analysis showed the spectro-spatial features of the mesolimbic network discriminated positive affective behaviors from affectless behaviors significantly better than chance in 10/10 subjects (Fig 2-C) and negative affective behaviors from affectless behaviors in 5/5 subjects (Fig 2-D). Group level results from all subjects replicated the successful performance of the positive and negative decoders found at the individual level (Mean ± sem AUC=**0.90**± 0.02, n= 10, Wilcoxon ranksum test, p < 0.0001 & **0.80**± 0.04, n= 5, p = 0.0012, Figure 2-E). Similar findings were also obtained using accuracy measures (number of true predicted samples / all samples) from all subjects (Figure 2-F and 2-G). A comparison of decoding performance in the positive and negative affective behaviors revealed that the positive decoders performed significantly better than negative decoders (Wilcoxon ranksum test, p = 0.04).

### Naturalistic affective behaviors were associated with increased activity in high frequency bands and decreased activity in low frequency bands

Our analyses thus far indicated that the RF decoders could discriminate positive and negative affective behaviors from affectless behaviors. We next investigated whether specific changes in spectral power across the mesolimbic network characterized each type of affective behavior, thus enabling the decoder models to distinguish them from behaviors lacking affect.

We used the trained decoder models to rank the spectro-spatial features that were best able to discriminate positive and negative affective behaviors from affectless behaviors in each subject. First, we ranked the importance of each spectro-spatial feature (herein, referred to as the “feature importance”), which was the amount of prediction error that each feature contributed to the RF decoding model (i.e. the amount of error when that feature was excluded from the decoder model) for each subject (Fig. 3-A shows the ranked features from positive decoders for an example subject, see Appendix). Next, we assigned an objective threshold (i.e. knee of the curve)(45) to the cumulative summation curve of the feature importance (Fig. 3-A inset, refer to methods) to select the neural features that were the dominant contributors to the positive or negative decoders for each subject.

**Figure 3:**
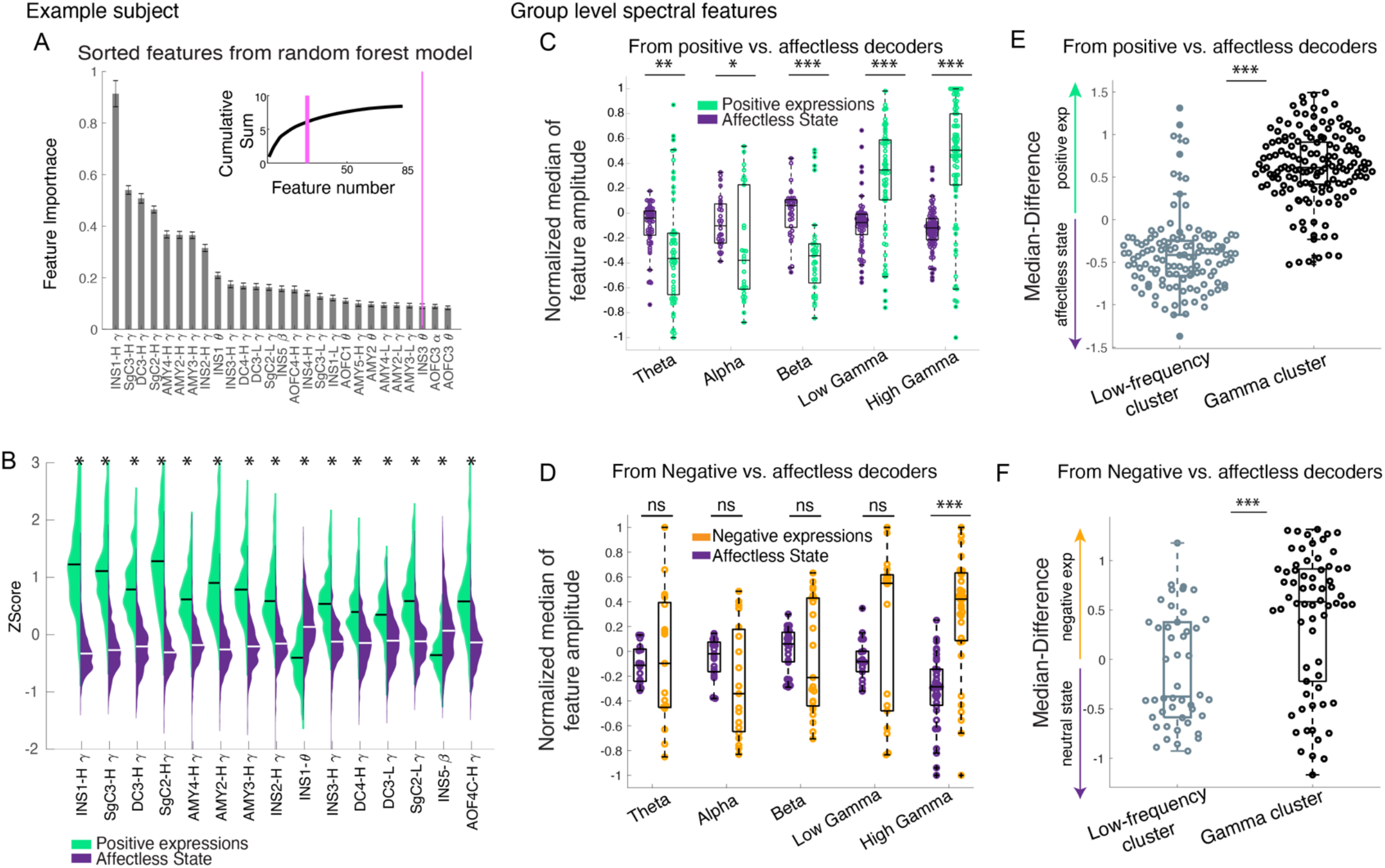
Gamma band (low and high) activity is increased during affective behaviors compared to affectless behaviors. A) The feature importance from the positive decoders is shown for the example subject (subject 1). Error bars are SEM across 100 datasets from the positive decoder models. The inset shows the cumulative summation curve of the feature importance; the pink vertical lines are the objective threshold that was used to select the top features. B) shows the sample distributions of the top 15 selected features for the positive affective behaviors (green) and affectless behaviors (purple). All sample distributions are significantly different from each other using non-parametric ranksum test. The median values of the gamma features are larger for the positive affective behaviors than the affectless behaviors while theta (INS1-theta) shows the inverse effect. C) The normalized median distributions of the positive affective behaviors and the affectless behaviors are shown for selected features across subject population. The median values from the positive decoders were first normalized to the maximum absolute spectral amplitude across selected features at a within-subject level and then pooled across all subjects (n = 10). In all frequency bands, the median values from the positive affective behaviors were significantly different from the affectless behaviors using a ranksum test: Theta p = 0.0001; Alpha p = 0.01; Beta, Low gamma, and High gamma p <0.0001. D) The normalized median distributions of negative affective behaviors and affectless behaviors are shown for selected features (n = 5). The median values were significantly different within the high gamma band only, p <0.0001. E & F) The median difference of the gamma cluster was selective to positive and negative affective behaviors and was significantly different from the low frequency cluster, respectively, p <0.0001. INS: insula, SgC = Subgenual cingulate, DC = dorsal cingulate, AMY: amygdala. H = high, L = low.

Using this unbiased feature selection approach, we observed that high gamma activity across multiple structures within the mesolimbic network (e.g., anterior insula, ventral ACC, dorsal ACC, and amygdala in the example subject) and low frequency activity (e.g., theta band in the anterior insula and amygdala in the example subject) were important features that contributed to the successful decoding of positive affective behaviors from affectless behaviors (Fig. 3-A depicts the pattern in one example subject). A similar spectral pattern emerged when we examined the features that characterized negative affective behaviors compared to affectless behaviors (Appendix Fig. S3-A).

After selecting the spectro-spatial features from each decoder type for each subject (Appendix, Fig. S4 & Fig. S5), we investigated the preference of these features for the affective behaviors. First, we extracted the z-score of sample distributions of these features in each subject (in the order of feature importance, as determined by the decoder models) for each behavior (Fig. 3-B depicts the example subject) and obtained the median amplitude of each. We next performed group level analyses using the selected spectro-spatial features to examine the extent to which the spectral patterns we observed within each subject held across individuals. To avoid bias toward subjects with stronger neural oscillatory activity, we normalized the median values by dividing them by the maximum absolute amplitude across all of the selected features in that subject and then grouped these values by their frequency bands. Across the sample, positive affective behaviors were characterized by increased power in the high and low gamma bands and decreased power in low frequency bands (e.g., theta, alpha, and beta) compared to affectless behaviors (Fig. 3-C). Like positive affective behaviors, negative affective behaviors were also characterized by increases in high gamma band activity and decreases in low frequency band activity (Fig. 3-D), but only high gamma power was significantly different between negative affective and affectless behaviors.

We performed an additional test of these results by conducting hierarchical clustering in each subject, which was an objective way to map the features that characterized the positive and negative affective behaviors from each decoder. While we observed common changes in spectral power across regions, clustering the features allowed us to control for possible collinearity between features (e.g., high gamma from multiple brain structures was increased during positive affective behaviors and might be driving our previous results, Fig. 3-A).

This clustering analysis identified two clusters—a “gamma” cluster and a “low frequency” cluster (Fig. S6 See Appendix for cluster naming)—from the positive and negative decoders that separated the positive, negative, and affectless behaviors based on spectral bands rather than regions (Appendix Fig. S3-B &C). These results suggested that, at an individual level, distributed spectral changes (i.e., simultaneous increases in gamma activity and decreases in low frequency activity in multiple regions) across the mesolimbic network characterized both positive and negative affective behaviors when compared to affectless behaviors (Appendix, Fig. S4 & Fig. S5). Although there were some exceptions to this pattern when examined at the individual level (e.g., for the positive decoder in subject 5, the clustering algorithm assigned a group to all spectral bands in OFC, which suggested in that case that a region was more important than the frequency band for distinguishing positive affective behaviors from affectless behaviors), in general, affective behaviors were separated from affectless behaviors along the spectral axis rather than along a spatial distribution (in which specific regions, not frequency bands, had primary roles in certain behaviors but not others) in the subjects when examined one by one.

We next investigated whether the spectral specificity for affective behaviors was observed across subjects, regardless of heterogenous spatial coverage. We computed a difference score for the selected feature within each cluster by subtracting the median activity(z-score) in that frequency band during the affectless state from the median activity during the positive or negative affective behaviors. This analysis found results consistent with those from the clustering analyses that were conducted at the individual subject level: compared to affectless behaviors, positive affective behaviors were characterized by higher median values in the gamma cluster (Fig. 3-E) and lower median values in the low frequency cluster (median of gamma cluster= 0.613 vs low frequency cluster = −0.416, ranksum test, p<0.0001). A similar pattern was found for negative affective behaviors when they were compared to affectless behaviors (Fig. 3-F, median of gamma cluster= 0.58 vs low frequency cluster = −0.37, ranksum test, p<0.0001). In sum, simultaneous increases in high frequency activity with low frequency desynchronization within the emotion network may serve as a global biomarker that is selective to affective behaviors.

### Despite common spectral changes across the emotion network during affective behaviors, distinct spatial patterns within the network distinguished between those of positive and negative valence

Our results revealed a common spectral pattern—increased high frequency activity and decreased low frequency activity—across the emotion network during affective behaviors (Appendix, Fig. S4 & Fig. S5). We next asked whether, despite this distributed network-level spectral pattern, certain brain regions within the network were more important to affective behaviors than other regions and whether certain regions played predominant roles in positive or negative affective behaviors.

We first conducted within-subject analyses to examine the extent to which high and low frequency spectral features were expressed in each region of the emotion network during positive and negative affective behaviors. Using the median difference scores for the gamma and low frequency cluster values (see previous section), scaled by their feature importance from the decoder models, we found that changes in both clusters were present across the emotion network during both types of affective behavior, consistent with our prior results (Fig 4-A & 4-C). These results again suggested that each region within the network participated in affective behaviors.

**Figure 4:**
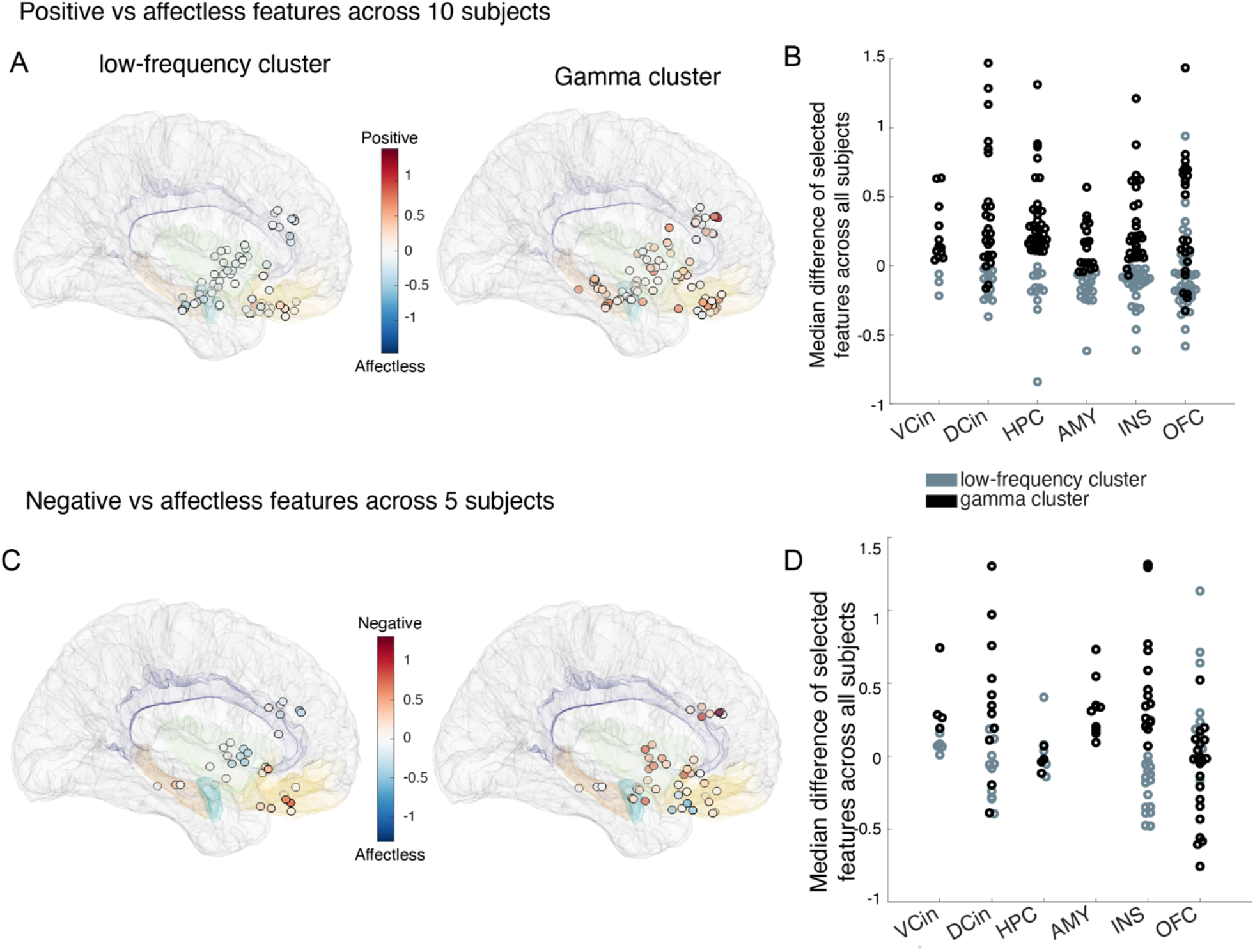
Both the “gamma” and “low frequency” clusters belong to a distributed network. As described in Figure 3, sample medians (z-score) for selected features for the positive and negative affective behaviors were normalized within each subject to account for between subject variability. For the gamma and low frequency clusters, the median features from the affectless behaviors were then subtracted from the medians of the positive and negative affective behaviors. The median differences were then scaled by their corresponding feature importance for each decoder type. Color bars in A and C represent this metric. A) Each of the electrodes are illustrated on the MNI brain with their corresponding contribution to the “low frequency” (left) and “gamma” (right) clusters from the positive decoders in all 10 subjects. White circles on the MNI brain indicate that the low frequency cluster is not as important as the gamma cluster, since it is scaled close to 0 value. B) The defined metric (see previous paragraph) from the gamma and low frequency clusters from the positive decoders are shown grouped by location. C& D) Similar as in A & B, pooled from negative decoders in 5 subjects. INS: insula, VCin = Ventral cingulate, DCin = dorsal cingulate, AMY: amygdala, OFC = orbitofrontal cortex, HPC = hippocampus.

We next conducted a more in-depth analysis to investigate whether certain regions across the network made stronger contributions to these cluster-level results. Analysis of the gamma and low frequency cluster activity in each region indicated that spectral changes in some structures within the emotion network were more important than others to affective behaviors. Compared to affectless behaviors, positive affective behaviors were characterized by increased gamma activity that was most prominent in ventral ACC and hippocampus, followed by dorsal ACC. Although gamma activity during positive affective behaviors was also elevated in other regions (i.e., anterior insula, amygdala, and OFC), the spectral pattern in these regions was more complex as both increased gamma activity and decreased low frequency activity contributed to positive affective behaviors (Fig. 4A-B & Fig. S6-E). When we compared negative affective behaviors to affectless behaviors, increased gamma activity in the amygdala was the most notable distinguishing feature (Fig. 4-C&D), and low frequency activity in the amygdala did not contribute to negative affective behaviors. The spectral changes in other regions (i.e., anterior insula, hippocampus, ventral ACC, and dorsal ACC) during negative affective behaviors were in the expected directions in both clusters (i.e., increases in gamma and decreases in low frequency activity, Fig. S6-E).

Our results indicated that certain regions within the emotion network contributed more strongly to different types of affective behavior when neural activity was quantified within different frequency bands. As a more rigorous test of this result, we next examined whether binary decoders could discriminate positive and negative affective behaviors from affectless behaviors when all spectral features were included. Here, we re-trained the within-subject positive (Appendix Fig. S7) and negative (Appendix Fig. S8) decoders in each region, one at a time. For each region, we included all of the spectral features from all of the depth electrodes implanted in that structure (and discarded the spectral information from all other regions across the network). Using a non-parametric Kruskal-Wallis test with Bonferroni adjustments for multiple comparisons applied on the AUC metric, we identified the top regions in each subject that could distinguish positive or negative affective behaviors from affectless behaviors significantly better than other regions (SI-Appendix Table S5 and S6).

To aggregate the within-subject decoding results across the group, we computed a ratio that reflected the number of subjects in whom each region was a significant predictor of positive or negative affective behavior (after correction for multiple comparisons) divided by the number of subjects in whom that region was sampled. Across the sample, widespread spectral changes in the anterior insula (7/9 subjects), amygdala (5/5 subjects), hippocampus (6/7 subjects), and ventral ACC (4/4 subjects) were more likely (>50% of subjects) to discriminate positive affective behaviors from affectless behaviors than spectral changes in the OFC (4/9 subjects) and dorsal ACC (4/10 subjects, Appendix Fig. S7 & Table S5). Spectral changes in the anterior insula (4/4 subjects), amygdala (1/1 subject), hippocampus (2/2 subjects), and dorsal ACC (3/5) were more likely than spectral changes in ventral ACC (1/2 subjects) or OFC (1/5 subjects) to distinguish negative affective behaviors from affectless behaviors (Appendix, Table S6 and Fig.S8).

Taken together, our results suggested affective behaviors are characterized by distributed spectral changes across the emotion network. Further examination of the relative contributions of different regions within the network indicated that some structures played more consistent roles in affective behaviors than others. The decoders found the anterior insula, amygdala and hippocampus were important contributors to affective behaviors in general, but the role of OFC was less consistent. Although the ACC was involved in both types of affective behaviors, there was also evidence that different ACC subregions played predominant roles in different types of affective behaviors. Whereas the ventral ACC (more than the dorsal ACC) contributed most dominantly to positive affective behaviors, the dorsal ACC (more than the ventral ACC) played a more prominent role in negative affective behaviors. Moreover, increased gamma activity in ventral ACC and hippocampus was a dominant contributor compared to low frequency bands in detecting positive affective behaviors. In one subject, gamma activity in the amygdala also played a dominant role in the cluster-level gamma increases that characterized negative affective behaviors. In sum, despite a common spectral pattern (increases in gamma and decreases in low frequency activity) across the network during both positive and negative affective behaviors, these results suggested there were distinct spectro-spatial features that characterized affective behaviors of differing valence.

### Three-state decoders demonstrated that high gamma activity across the emotion network distinguished among naturalistic behaviors

Our previous analyses using binary decoders showed we could decode positive and negative affective behaviors from affectless behaviors and identify the spectro-spatial features that characterized each behavior type. We next compared these three types of behavior directly by training within-subject multi-class RF decoders to distinguish among positive, negative, and affectless behaviors in three subjects who displayed sufficient instances (i.e., >=15 samples, within each portion, or fold, of the dataset). Using all of the spectro-spatial features from the emotion network, the multi-class decoder distinguished all three affective behaviors with an average accuracy of 0.68 ± 0.016, which was significantly above chance level (33%) in each of the subjects (Fig. 5-A, Appendix Fig. S10 &Table S7).

**Figure 5.**
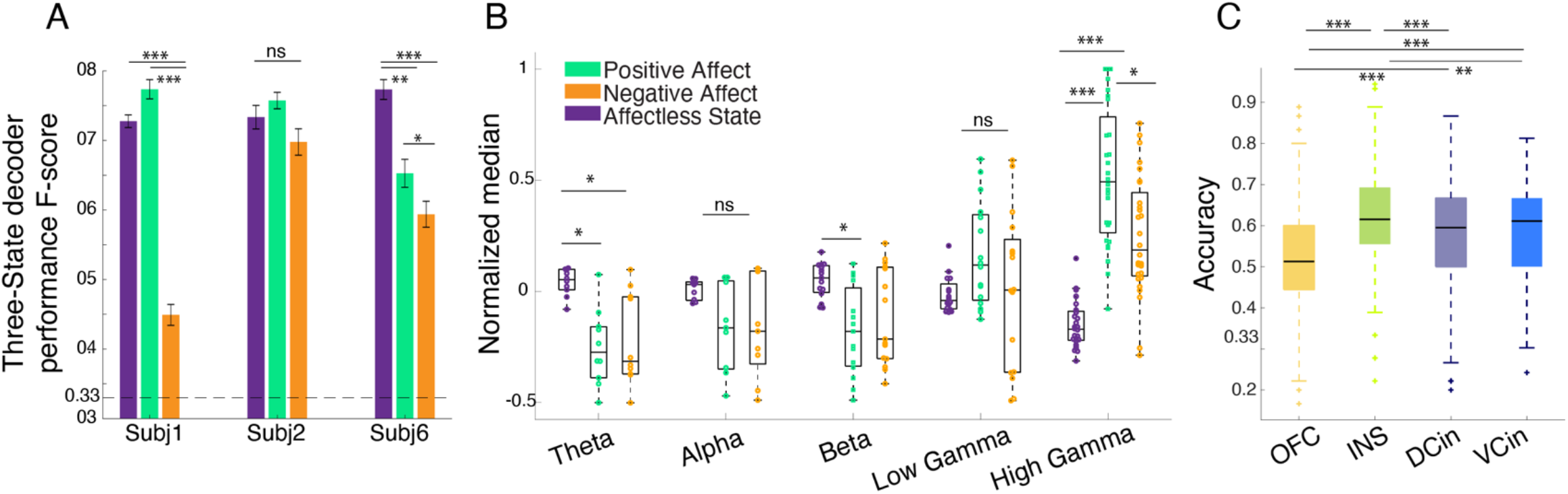
The three-state decoder distinguished among positive, negative, and affectless behaviors using the spectro-spatial features of the emotion network. A) F1-scores for the three-class RF models from the three subjects. All F1-Scores are significantly above chance level (33%, dahsed lines) and different from the shuffled models, pairwise ranksum test). Asterisks represents multiple comparison Kruskal Wallis test corrected with Bonferroni method across the F1-scores of each affective behavior within each subject; in subject 1 & 6 both positive and affectless behaviors have significantly larger performance than negative behaviors. In subject 6 positive behaviors are classified with larger performance compare to negative ones with p= 0.047. B) The median distribution of the selected features across the three subjects. multiple comparison test shows: Theta: positive and negative affective behaviors differed from affectless behavior, p = 0.0034 and p = 0.01, respectively. Alpha and Low Gamma: no significant differences were observed. Beta: only positive affective behavior differed from affectless behavior, p = 0.01. High gamma: positive and negative affective behaviors differed from affectless behavior p <0.0001). negative and positive behaviors differed from each other, p = 0.047. C) The three-state decoder models were trained using the spectral features from each region and then pooled across the three subjects. connotations as in figure 4. OFC is from 4 probes implanted in 3 subjects, INS and DCin are from 3 probes from 3 subjects, and VCin 2 probes from 2 subjects. Statistics regarding panel C using a corrected multiple comparisons test is as following: the anterior insula was significantly different from dorsal ACC and OFC with p <0.0001 and from ventral ACC with p <0.001. Ventral and dorsal ACC both are different from OFC with p <0.0001. All *** means p <0.0001 and All ** means p <0.001

The multi-class decoding performance was better for positive than negative affective behaviors in each of the three subjects (mean+/-std F1-scores= 2*(precision*recall)/(precision+recall); 0.74 ± 0.013 for affectless behaviors, 0.72 ± 0.037 for positive behaviors, and 0.57 ± 0.07 for negative behaviors). This result, though surprising, was consistent with our prior finding with the binary decoders, which distinguished positive affective behaviors from affectless behaviors more effectively than negative affective behaviors from affectless behaviors (Fig. 2-E). The decoding performance of positive versus negative affective behaviors was consistent across subjects, suggesting these results were robust and not a reflection of larger number of instances of positive than negative affective behaviors included in each analysis (Appendix, Fig. S9).

We next looked across the group to examine which features were most important for decoding the three behaviors with the multi-class decoder across subjects. Group-level analyses of selected spectro-spatial features from the three subjects (grouped by spectral band) demonstrated that high gamma activity was greater during positive and negative affective behaviors than during affectless behaviors and that high gamma activity discriminated among these three behavior types (Fig. 5B). Low frequency activity in the theta and beta frequency bands, in contrast, was decreased during both positive and negative affective behaviors compared to affectless behaviors but did not significantly differ between affective behaviors of differing valence. These findings suggested that while increased high gamma and decreased lower frequency power across the emotion network characterized affective behaviors in general, only high gamma activity discriminated positive from negative affective behaviors. Although we could not determine whether the different levels of high gamma activity during positive and negative behaviors were due to differences in the valence or the intensity of these states, this result suggested that, despite common spectral changes during moments of affect, the different frequency bands also made distinct contributions to various types of affective behavior.

To investigate whether spatially localized activity within the emotion network could differentiate the three behaviors, we trained the three-class decoders in each subject using the spectral features from each region, one at a time, and then compared the performances of the resulting models. We found that multiple regions successfully decoded the affective behaviors in each subject with accuracy (number of true predicted samples / all samples) significantly above chance (33%; Appendix Fig. S11). The anterior insula (3/3 subjects) and dorsal ACC (2/3 subjects), in particular, were important for distinguishing among positive, negative, and affectless behaviors followed by the amygdala (1/1 subject), hippocampus (1/1 subject), and ventral ACC (1/2 subjects).

To examine these results in more detail, we concatenated the decoder accuracies from each subject in regions that were sampled in at least two subjects (i.e., the amygdala and hippocampus were not included here because they were each sampled in one subject): anterior insula (3/3 subjects), ventral ACC (2/3 subjects), dorsal ACC (3/3 subjects), and OFC (3/3 subjects). Using a Kruskal-Wallis test with Bonferroni-corrected tests for multiple comparisons, we found that the accuracy of the anterior insula was the highest (mean ± sem = 0.62 ± 0.006) followed by the ventral ACC (0.58 ± 0.007), dorsal ACC (0.58 ± 0.008), and, lastly, the OFC (0.52 ± 0.006) in distinguishing behaviors with the three-state decoder (Fig. 5-C). Indeed, the level of high gamma power, as determined by the median power from the anterior insula, ventral ACC, and dorsal ACC channels showed stratification across the three behaviors (Appendix, Fig. S12). We additionally observed that high gamma features had significantly larger importance values than the low frequency features in the decoder models that were trained on all spectro-spatial features (i.e., full models, Appendix, Fig. S13). There was no significant difference between high gamma and low frequencies in regard to specific regions.

## Discussion

We found evidence that direct neural recordings of the human emotion network could discriminate naturalistic affective behaviors from affectless behaviors with high accuracy. We trained machine learning decision tree-based models on the spectro-spatial features of the emotion network and successfully decoded positive affective behaviors (with up to 93% accuracy) and negative affective behaviors (with up to 78% accuracy) from affectless behaviors using binary decoders in individual subjects. Investigations of the spectro-spatial features that characterized affective behaviors across subjects revealed some features that were characteristics of both positive and negative affective behaviors and others that were specific to each. In general, affective behaviors were associated with coordinated changes across the emotion network including increased activity in high frequency bands (i.e., gamma band) and decreased activity in low frequency bands (i.e., theta, alpha, and beta bands). When we examined the contributions of different structures within the network to these overarching spectral patterns, certain regions emerged as playing more central and consistent roles in affective behaviors (i.e., anterior insula, amygdala, and hippocampus) than others (i.e., OFC). Although there were spectro-spatial similarities between positive and negative affective behaviors, different spatial topographies also characterized each. The ACC also played a central role in affective behaviors, and activity in different subregions discriminated between affective behaviors of differing valences: whereas the ventral ACC contributed more strongly to positive affective behaviors, the dorsal ACC contributed more strongly to negative affective behaviors. In a subset of subjects, a multi-class decoder that compared all three behaviors directly highlighted that high gamma activity across the emotion network was increased during affective behaviors compared to affectless behaviors and that these high gamma increases were more substantial during positive than during negative affective behaviors. This analysis also emphasized that the anterior insula and ACC played more central roles in affective behaviors than other regions, such as OFC.

Consistent with previous studies of emotion, our results indicate a distributed network supports emotions and the affective behaviors that accompany them. Although fMRI studies cannot measure neural activity in different frequency bands, noninvasive EEG studies have found consistent increases in gamma band activity in the emotion network during emotions (9, 28, 46), the patterns in lower frequency bands are less clear. Studies that have investigated changes in lower frequency bands during emotions have found more variable results including decreased beta band activity in ACC in response to negative stimuli (30); increased theta band activity in frontal cortex during negative and positive emotions including anger, disgust, and joy (24, 47); and increased overall theta band activity during positive emotions such as amusement and joy (35).

Despite numerous methodological differences between prior studies and the approach we took here in studying naturalistic affective behaviors, we also found consistent increases in gamma band activity during both positive and negative affective behaviors that subjects spontaneously expressed throughout their hospital stays. In general, we found simultaneous increases in high frequency activity and decreases in low frequency activity across the emotion network during both types of affective behaviors compared to affectless behaviors. Our results are consistent with a previous neurostimulation study that demonstrated that transient increases in gamma power and decreases in low frequency power also accompany and promote cognitive functioning (48). Although many unanswered questions remain regarding the role of lower frequency bands in emotion, our results suggest affective behaviors that arise in ecologically valid contexts may engage similar neural mechanisms—particularly when measured in the high frequency bands— as those observed during more controlled experimental tasks. Taken together, the results of the spectral analyses indicated that both positive and negative affective behaviors were characterized by increased gamma band activity and decreased low frequency activity across the emotion network.

An examination of the spatial topography of emotion network activity revealed that some regions contributed more strongly than others to the distributed spectral changes that characterized affective behaviors. While some previous studies have emphasized the role of the anterior insula in negative emotions, namely disgust (6, 23), the binary and multi-class decoders pointed to a central role for the anterior insula in both positive and negative affective behaviors. The anterior insula, a key hub in interoceptive pathways(13, 19) that support awareness of internal feeling states, was a key contributor to both types of behavior. The amygdala and hippocampus were also important for decoding both positive and negative affective behaviors from affectless behaviors. The amygdala is critical for generating emotions and, though often associated with negative emotions (9, 21,24, 31,49), activates during negative and positive states of sufficient intensity and has been implicated in networks supporting affiliative behavior as well as threat responding (33, 50). Deep brain stimulation studies have also found that brief perturbation of amygdala subregions can induce rapid negative (31) and positive affective reactions (ERP at ~200-400 ms) (24). In one subject with amygdala coverage, we found gamma activity in the amygdala played a prominent role in negative affective behaviors, but more evidence is needed to corroborate this result. The hippocampus has dense structural connections with the amygdala (51, 52) and is essential for emotional memories, which may be relived during spontaneous moments of affect. Although the role of the hippocampus in mood and emotion is still debated, recent iEEG studies have found that lower mood is associated with greater beta coherence between amygdala and hippocampus (36), which suggests both structures, and their interaction, may be critical for both positive and negative affective states.

Despite some common spectro-spatial patterns during positive and negative affective behaviors, different regions within the emotion network also made distinct contributions to each type of behavior. The ACC, in general, was important for affective behaviors, and increased gamma activity in ACC characterized both positive and negative affective behaviors. Within the ACC, however, differences emerged, in the degree to which its subregions participated in positive and negative affective behaviors. Our results suggested ventral ACC was more critical for positive affective behaviors than dorsal ACC and that dorsal ACC was more critical for negative affective behaviors than ventral ACC. The ACC is comprised of several cytoarchitectonically distinct subregions, and these findings are consistent with neuromodulation studies that have shown that whereas stimulation of ventral ACC can cause laughter and mirth (18, 53), stimulation of dorsal ACC can cause feelings of doom and perseverance. The dorsal and ventral ACC have different anatomical projections to autonomic and motor centers that are critical for emotions(54), and our results suggest ACC subregions engage these distinct pathways to produce positive and negative affective behaviors.

Relative to other regions in the emotion network, the OFC played a less consistent role in affective behaviors. The OFC, especially in lateral areas, is critical for emotion regulation, cognitive control, and behavioral inhibition (55–57). One recent neuromodulation study found that stimulation of lateral OFC decreased theta activity across the emotion network and improved mood (37), a finding that suggested that suppression of low frequency activity yielded emotional benefits. Although our results indicated that activity in low frequency bands (e.g., theta, alpha and beta) across the emotion network decreased during both positive as well as negative affective behaviors, OFC was not robustly selected in either type of affective behavior. Our findings suggest the OFC may be engaged in different ways depending on the emotional context, thus making its contribution to affective behaviors more variable across instances and subjects.

The present study offers a novel window into the neural mechanisms of the emotion network. In the field of affective neuroscience, there is ongoing debate regarding the degree to which different emotions have unique or shared representations in the brain. Although some suggest the emotion network activates in the same way during even very different affective states, others emphasize the biological uniqueness of each emotion and posit there are dissociable—albeit difficult to discern—patterns of neural activity that accompany them. Our results help to integrate these contrasting views by beginning to elucidate how a distributed emotion network might produce different affective states. Although positive and negative affective behaviors differ in valence, both can vary in arousal or intensity levels. Some of our results suggested that common changes in emotion network activity occurred during affective behaviors in general, regardless of whether the behaviors were positive or negative. Across the network, for example, increases in gamma activity and decreases in low frequency activity characterized both positive and negative affective behaviors. There were also regions (i.e., anterior insula, amygdala, hippocampus, and ACC) that contributed more strongly than other regions (i.e., OFC) to both types of behavior. We speculate that the shared gamma activity in these regions during both positive and negative affective behaviors may reflect the role these regions played in representing arousal or the intensity of emotional experience, a dimension of affect that may have been on a comparable scale during both types of behavior. Our results also indicated, however, that different structures within the emotion network also made distinct contributions to positive and negative affective behaviors and may have helped to shape these distinct affective states. Whereas gamma band activity in the ventral ACC, hippocampus and dorsal ACC contributed more to positive affective behaviors, increased gamma activity in the amygdala in one subject played a prominent role in negative affective behaviors while no low frequency in these regions were prominent contributors to decoding these behaviors (Fig 4D & E). A distributed emotion network that activates through a combination of spectral changes (that are common to affective behaviors in general) and spatial changes (that are specific to positive or negative affective behaviors in particular) would be a flexible system that is well-equipped to produce a variety of affective states.

The present study has several limitations to consider. First, we analyzed neural activity during positive and negative affective behaviors, valence-based behavioral categories that included a variety of different activities. Although the positive affective behaviors were fairly uniform (mostly comprised of smiling and laughing), the negative affective behaviors were more heterogeneous and included a range of unpleasant including expressions of pain, frustration, or low mood. Our results suggested it was more difficult to decode negative than positive affective behaviors (Fig 2 and Fig. 5), which was likely due, in part, to the greater variability among the behaviors that comprised the negative category. Moreover, the larger decoding performance was robust to larger sample number of positive affective behaviors compared to negative affective behaviors (Appendix, Fig S9). Additional studies are needed to determine how the emotion network produces behaviors that characterize distinct types of negative as well as positive emotions and whether differences in emotion network activity emerge when examined during specific emotions.

Second, due to the unconstrained nature of our study, we did not have measures of self-reported experience and, thus, were unable to make direct comparisons between positive and negative affective behaviors of comparable intensities. Our results indicated there were many similarities between positive and negative affective behaviors at the neural level. Given that positive and negative affective behaviors, by definition, differ in valence, we speculate that the common neural activity patterns that characterized both positive and negative affective behaviors compared to affectless moments may have reflected arousal (or emotional experience intensity) that accompanied during behaviors of either valence. Future studies that measure self-reported experience, an accessible metric of internal feelings (58, 59), as well as behavior will be needed to shed light on this question and to compare positive and negative emotions of differing intensities.

Third, electrode placement was based on clinical needs and, thus, coverage of the emotion network varied somewhat across subjects. Even within a single region, electrodes may have sampled distinct subregions in different subjects, which may have increased the functional variability across subjects. There was also variability in the number and types of affective behavior that subjects expressed spontaneously throughout their hospital stay. We leveraged the power of our longitudinal dataset by training subject-specific classifiers to decode affective from affectless behaviors and then looking across the subjects to identify results that were robust across the sample. Although iEEG, unlike fMRI or EEG, has access to some deep structures, it was still not possible to sample other structures within the emotion network (e.g., periaqueductal gray).

In summary, we used state-of-the-art machine learning approaches (60) to decode naturalistic affective behaviors from direct recordings of the human emotion network. Measurement of neural activity in different frequency bands in multiple regions across the emotion network uncovered new information about the spectro-spatial features that characterize affective behaviors. To further improve the interpretation of the identified spectro-spatial features by the random forest model, we employed objective threshold selection followed by hierarchical clustering. We additionally trained support vector machine (SVM) classifiers using the selected feature set that were derived from the RF models. The resulting SVM models reached similar performance as RF models (Appendix Fig. S14) confirming the robustness of this feature selection method.

The combination of methods that we utilized here provides a potential strategy to ask whether various affective states are dissociable at the neural level and to identify their neuronal characteristics from large datasets. Complex, real-time decoding models trained on neural activity in sensorimotor and language cortices have made it possible to design brain-computer interfaces for those who suffer from limb (61) or speech malfunctions (38), but similar advances are lacking in neuropsychiatry, and it remains difficult to relate neural signals to the complexity of emotions and mood (62). More sophisticated neuroanatomical models of affective symptoms would inform personalized treatments for neuropsychiatric and affective disorders and would help to identify biomarkers that could be monitored in treatments such as closed-loop based neurostimulation.

## Materials and Methods

### Subjects & Inclusion criteria

Subjects included 11 patients (6 females, 5 males, age: 20-43, Table S2), who has been diagnosed with treatment-resistant epilepsy and were undergoing (iEEG) implantation for seizure localization. Subjects that accepted to provide informed consent, or at least had implanted electrodes in 3/5 regions, or the number of behavioral emotional expressions was sufficient enough to train the RF classifiers were included in the study (Table S1 and Table S3). All procedures were approved by the University of California, San Francisco Institutional Review Board. All subjects gave written informed consent to participate in the study prior to surgery.

### iEEG and Behavioral Data Acquisition

Over multiple days of monitoring, subjects underwent continuous 24-hour audio, video recording and iEEG monitoring through the Natus clinical recording system as a part of routine clinical care. Electrophysiological data were collected at sampling rates at either 512 Hz or 1024 Hz. All mesolimbic structures were sampled by subdural grid, Ad-Tech 4-contact strip and Ad-Tech 4/10- contact depth electrodes (10mm or 6mm center to center spacing). 2 patients had mini grids implanted on OFC.

### Behavioral Annotations

Trained human raters manually annotated these recordings during instances of behavioral and emotional expression. The affective behaviors of the subjects that were of particular interest for the purpose of this study and were coded in ELAN as a tick at the timepoint (on a millisecond basis) where the emotional expressions occurred. These instances were classified as behaviors the patient exhibited either through visual examination of the video or through conversations the patients had with others in their hospital room. Affective behaviors consisted smiling, laughing, positive verbalization, negative verbalization, or the physical display of pain.

Detailed materials and methods regarding behavioral annotations, iEEG time-frequency and decoder analyses are provided in the Appendix.

### Data and code availability Statement

Data used to generate the findings of this study will be available upon request to the Lead Contact.

## Acknowledgments

Authors would like to acknowledge Chang lab members, Ben Speidel, Dharshan Chandramohan, Kristin Sellers, Lowry Kirkby, and raters Nicole Goldberg-Boltz, Lucia Bederson, Muriel Solberg, Claire Eun, Joe Gordon, Dale Tager, Vivian Cheng, Nikhita Mummaneni, Nikhita Kunwar. This research was funded by the Defense Advanced Research Projects Agency (DARPA) under Cooperative Agreement Number W911NF-14-2-0043. The views, opinions, and/or findings contained in this material are those of the authors and should not be interpreted as representing the official views or policies of the Department of Defensei or the U.S. Government.

## Author Contributions

M.B. performed all analysis. M.D., D.L.W, and E.F.C. designed the study. D.L.W. and M.D. assisted with subject recruitment, data collection and leading behavioral annotations. M.B. and A.N.K. conceptualized the analytical framework. M.B., M.D. and A.S. performed neural data cleaning from epileptiform activity. M.B., A.N.K. and V.E.S. wrote the manuscript with input from other authors. H.E.D. and E.F.C. supervised the experimental work.

